# Photophysics-informed two-photon voltage imaging using FRET-opsin voltage indicators

**DOI:** 10.1101/2024.04.01.587540

**Authors:** F. Phil Brooks, Hunter C. Davis, Pojeong Park, Yitong Qi, Adam E. Cohen

## Abstract

Microbial rhodopsin-derived genetically encoded voltage indicators (GEVIs) are powerful tools for mapping bioelectrical dynamics in cell culture and in live animals. Förster resonance energy transfer (FRET)-opsin GEVIs use voltage-dependent changes in opsin absorption to modulate the fluorescence of an attached fluorophore, achieving high brightness, speed, and voltage sensitivity. However, the voltage sensitivity of most FRET-opsin GEVIs has been reported to decrease or vanish under two-photon (2P) excitation. Here we investigated the photophysics of the FRET-opsin GEVIs Voltron1 and 2. We found that the voltage sensitivity came from a photocycle intermediate, not from the opsin ground state. The voltage sensitivities of both GEVIs were nonlinear functions of illumination intensity; for Voltron1, the sensitivity reversed sign under low-intensity illumination. Using photocycle-optimized 2P illumination protocols, we demonstrate 2P voltage imaging with Voltron2 in barrel cortex of a live mouse. These results open the door to high-speed 2P voltage imaging of FRET-opsin GEVIs *in vivo*.

**Teaser:** Voltage sensitivity in FRET-opsin indicators comes from a photocycle intermediate, reachable via optimized 2P excitation.

## Introduction

Genetically encoded voltage indicators (GEVIs) are a powerful class of fluorescent probes for mapping bioelectrical signals.^1^ These tools have been used in multiple species^2–7^ and at levels of biological organization from sub-cellular^8–10^ to organ-wide^11–13^. Microbial rhodopsin-based GEVIs have fast (submillisecond) responses to voltage steps, and good voltage sensitivity.^14,15^ The first opsin-based GEVIs relied on the near infrared fluorescence of the retinal cofactor, but this signal was very dim.^5,16,17^ In fluorescence resonance energy transfer (FRET)-opsin GEVIs, voltage-dependent changes in the opsin absorption spectrum modulate the efficiency of FRET from an attached fluorophore, leading to modulation of the fluorophore fluorescence (Fig. 1a).^18,19^ This approach has been demonstrated with fusions of fluorescent proteins to microbial rhodopsins,^7,20^ and with fusions of the HaloTag receptor, which can be covalently loaded with a small-molecule organic dye.^2,21^ FRET-opsin GEVIs are fast, bright, and sensitive^18,21–23,2^.

**Figure 1:**
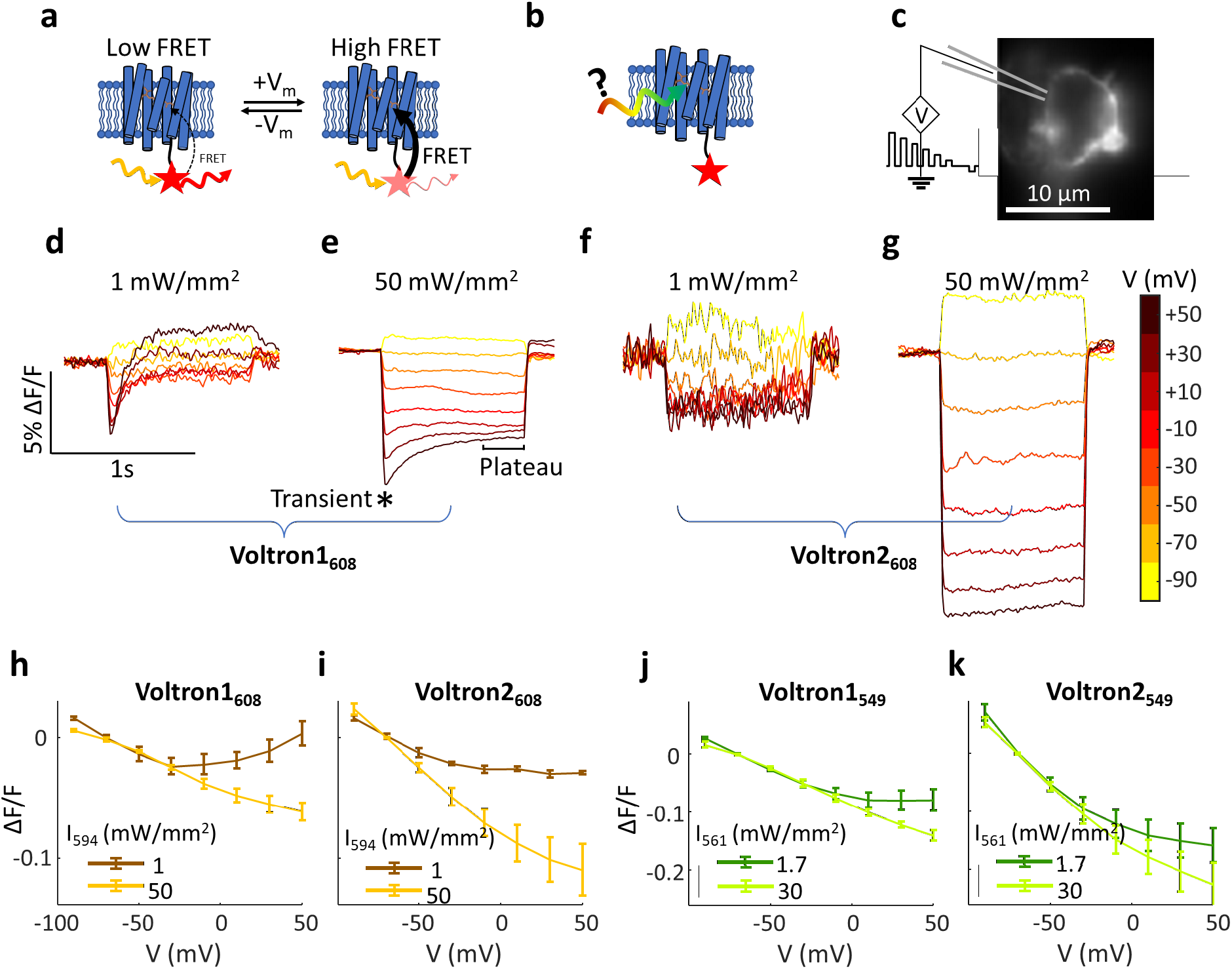
Illumination-dependent performance of FRET-opsin voltage indicators. a) A simple model of a FRET-opsin GEVI. A fluorescent FRET donor is optically excited and can relax either by fluorescence or by FRET to the retinal chromophore. Voltage-dependent shifts in the retinal absorption spectrum modulate the fluorescence of the donor. b) The light used to excite the FRET donor may also excite the retinal directly, driving photo-transitions in the opsin and changing the voltage-sensing properties of the GEVI. c) Fluorescence image of a HEK-293T cell expressing Voltron2_608_ and subject to voltage clamp. d,e) Voltage step responses from cells expressing Voltron1_608_ at low (1 mW/mm^2^) and high (50 mW/mm^2^) illumination intensities, λ = 594 nm. Transient and plateau phases of the response are indicated. At low intensity, steady state fluorescence responses showed a non-monotonic dependence on membrane voltage. f,g) Same as (d,e) for Voltron2_608_, with the same ΔF/F vertical scale. h-k) Plots of steady-state ΔF/F vs. V for (h) Voltron1_608_, (i) Voltron2_608_, (j) Voltron1_549_ and (k) Voltron2_549_. Each curve is plotted for dim (dark colors) and bright (light colors) illumination. For all reporter combinations, both the slope and the shape of the curve were sensitive to illumination intensity. Error bars represent s.e.m. from 3-6 cells.

A key challenge in voltage imaging is to resolve signals within light-scattering tissues, such as the brain. Most applications of voltage imaging to-date have used one-photon (1P) excitation. While structured illumination, far-red excitation, and use of photo-activatable GEVIs^24^ can partially reduce the background from scattered light, 1P voltage imaging is still limited to imaging the top ∼250 μm of brain tissue. Two-photon (2P) excitation has been transformative for calcium imaging *in vivo*, so there has been substantial interest in developing 2P voltage imaging systems.^25,26^ The FRET-opsin GEVIs would be attractive targets for 2P voltage imaging, but, for reasons that have remained mysterious, most FRET-opsin GEVIs show little or no voltage sensitivity under typical 2P illumination conditions, even when the fluorescence count-rate is high enough that voltage-induced fluorescence changes should be detectable.^27,28^ Furthermore, these same samples can return to showing voltage sensitivity under 1P illumination after 2P illumination.^27^ These observations led us to explore the photophysical basis of voltage sensitivity in FRET-opsin GEVIs.

In the simplest model of a FRET-opsin GEVI (Fig. 1a), a voltage-insensitive FRET donor is optically excited. The opsin FRET acceptor sits in a voltage-sensitive equilibrium between two states, one of which quenches the donor more efficiently than the other. This simple model produces two important predictions. First, the fractional response of the donor fluorescence to voltage (i.e. ΔF/F vs. V) should be a function only of voltage and not of any illumination parameters. Second, any excitation method (*e*.*g*., 1P or 2P) that produces the same donor excited state should produce the same voltage-sensitive fluorescence signal. The documented failure of 2P voltage imaging with FRET-opsin GEVIs suggests that this simple picture is inadequate.

The light used to excite the FRET donor can also interact with the opsin acceptor directly (Fig. 1b). Microbial rhodopsins have complex photocycles, with at least seven spectroscopically distinguishable states, and a variety of light- and voltage-modulated transitions.^29–34^ Indeed, in wild-type Archaerhodopsin 3, voltage-sensitive retinal fluorescence comes from a photocycle intermediate termed the “Q state”, and exciting this fluorescence requires sequential absorption of three photons.^32^ The engineering of QuasAr1, QuasAr2 and the Archon variants caused the voltage sensitivity to appear as a 1-photon process, but this signal still arises from a complex photocycle involving multiple absorbing states^34–36^.

The complexity of opsin photocycles has been harnessed to create light-gated voltage integrators,^37^ light-gated voltage sample- and-hold motifs,^37^ reporters of absolute voltage,^38^ and photo-activated voltage indicators.^24^ Thus we hypothesized that the voltage-sensing properties of FRET-opsin GEVIs might depend in a complex way on the intensity, wavelength(s), and time-course of illumination; and that an understanding of these dependencies might point to illumination protocols which enabled 2P voltage imaging.

## Results

We expressed Voltron1^21^ or Voltron2^2^ in HEK-293T cells and labeled the samples with HaloTag ligand dye JF_608_ (Methods). We selected this dye because it has been useful in all-optical electrophysiology experiments with Voltron1 and 2.^8,10^ We then used whole-cell voltage clamp to vary the membrane voltage and we recorded the fluorescence under continuous 594 nm 1P illumination at different intensities (Fig. 1c).

We first applied a series of voltage steps from a holding potential of −70 mV to voltages between −90 mV and +50 mV (Fig. 1d-g). At high illumination intensity (50 mW/mm^2^), both GEVIs showed an approximately linear and negative-going dependence of steady-state fluorescence on membrane voltage, with slopes ΔF/F = −0.054 ± 0.007 per 100 mV (Voltron1, n = 4 cells) and ΔF/F = −0.11 ± 0.02 per 100 mV (Voltron2, n = 5 cells), where ΔF was measured relative to F at V = −70 mV (Fig. 1h, i). For depolarizations to > 0 mV, Voltron1 showed an initial transient fluorescence peak (Fig. 1e), but Voltron2 did not (Fig. 1g). These data are all consistent with prior reports^2,21^.

At low illumination intensity (0.97 mW/mm^2^), the voltage responses of both GEVIs changed dramatically. The initial transient fluorescence response of Voltron1 maintained its approximately linear negativegoing dependence on voltage. However, the steady-state F vs. V response of Voltron1 became nonmonotonic. For small depolarizations relative to −70 mV, the fluorescence decreased, as at high intensity. But for depolarizations to > −20 mV, the Voltron1 fluorescence increased as voltage increased, and at voltages > +30 mV, the steady-state fluorescence was actually brighter than at V = −70 mV (Fig. 1d, h).

For Voltron2 at low illumination intensity, the voltage step-response maintained its top-hat structure (Fig. 1f), but the overall voltage sensitivity decreased nearly twofold for small depolarizations around −70 mV and the voltage response leveled off for voltages > −20 mV. Qualitatively similar intensity-dependent changes in the transient and steady-state voltage responses were observed when the two GEVIs were loaded with JF_549_ and excited at 561 nm (Fig. 1j, k).

To quantify the influence of illumination intensity on voltage sensitivity, we performed voltage-clamp experiments at 1P illumination intensities spanning nearly four orders of magnitude, from 0.06 mW/mm^2^ to 100 mW/mm^2^ (Fig. 2a,b). At each intensity, we clamped the voltage at −70 mV and then measured the fluorescence responses to a 100 mV depolarizing step to +30 mV. We plotted separately the initial and steady-state fluorescence responses, as marked in Fig 1e (for Voltron2, initial and steady state fluorescence were indistinguishable).

**Figure 2:**
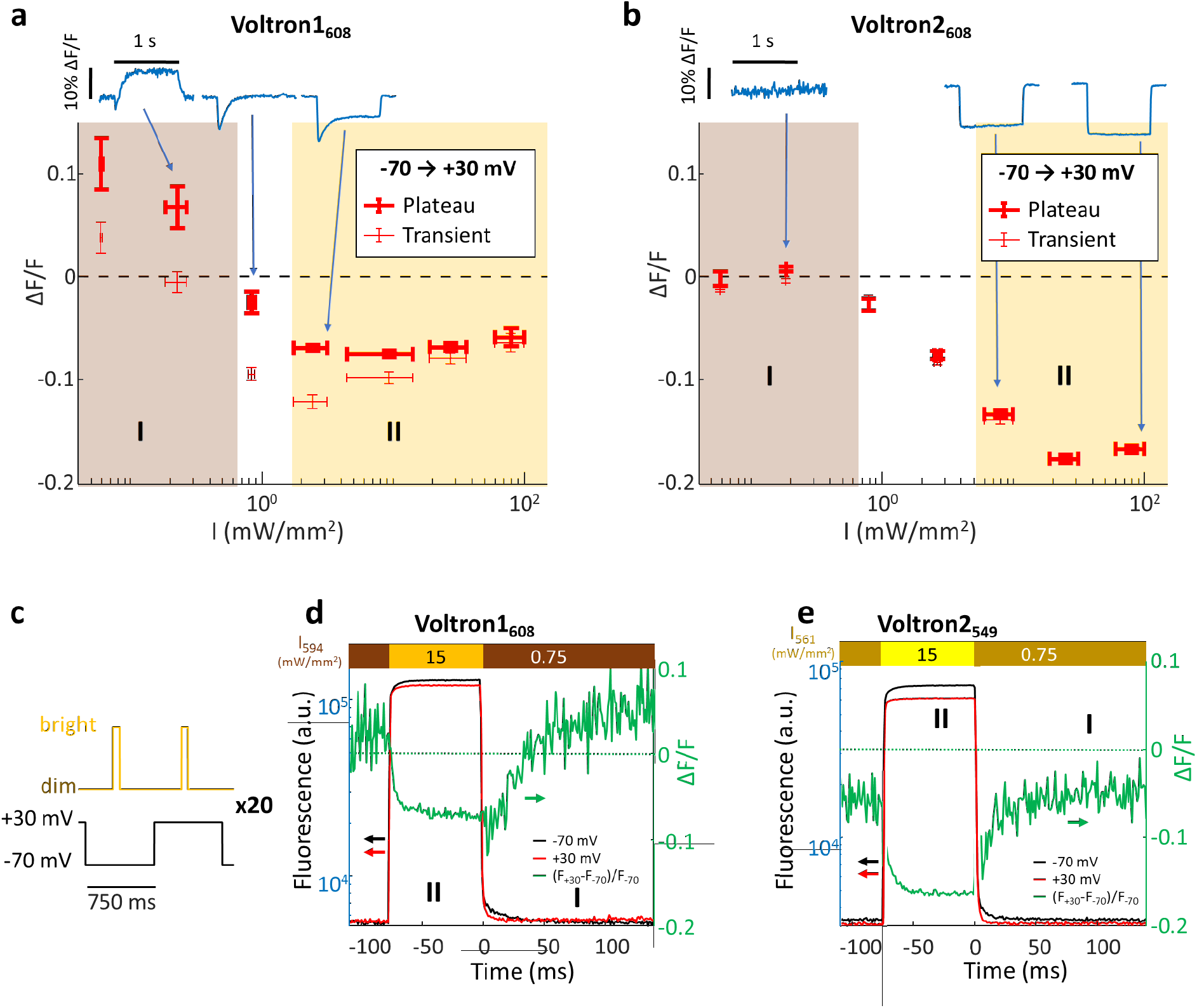
Intensity-dependent voltage sensitivity of Voltron GEVIs. a) Fractional sensitivity (ΔF/F) of Voltron1_608_ as a function of illumination intensity for a voltage step from −70 to +30 mV. Heavy markers denote the steady-state response, thin markers denote the initial transient response (n = 5 cells). Horizontal error bars denote the range of illumination intensities binned into one measurement; vertical error bars denote s.e.m. The traces in blue show representative step responses from dim, moderate, and bright illumination. b) Same as panel (a), but for Voltron2_608_. (n = 3 cells). c) Protocol for measuring dynamic GEVI responses to a change in illumination intensity. Dynamics in the “dark” were probed by very dim (0.75 mW/mm^2^) illumination; bright pulses (75 ms, 15 mW/mm^2^) transiently populated the voltage-sensitive stage. d) Fractional voltage sensitivity of Voltron1_608_ (green, right axis) was calculated from the difference between the fluorescence at −70 mV (black, left axis) and at +30 mV (red, left axis). Sensitivity emerged with a time constant of 5 ms, reached a plateau during the light pulse, and then declined with a time constant of 46 ms after the bright pulse. e) Same as (d) for Voltron2_549_. Sensitivity emerged with a time constant of 7 ms and declined with a time constant of 18 ms.

The data showed several surprising features. For an idealized GEVI, one would expect both ΔF and F to be proportional to intensity, and their ratio (ΔF/F) to be independent of intensity. We found that for both indicators, ΔF/F depended strongly on illumination intensity. This observation shows that statements of FRET-opsin voltage sensitivity are only meaningful if illumination intensity is specified. For both GEVIs, the voltage sensitivity was greatest (in absolute value), and the illumination intensity dependence leveled off around 10-30 mW/mm^2^. By good fortune, this intensity regime is typically used in neural recordings because it produces high enough per-cell count rates to observe neural dynamics over shot noise. This coincidence may explain why the low-intensity anomalous responses of these GEVIs were not previously reported.

These data also show the disparate effects of illumination intensity on different response timescales. At low illumination intensity, Voltron1 showed the unexpected inversion of sensitivity to 100 mV voltage steps (Fig. 2a). Further, at low intensity Voltron1 showed enhanced disparity between the transient and steady-state responses to 100 mV voltage steps. While the sensitivity of Voltron2 was in general superior to Voltron1, for illumination intensities between 1 – 10 mW/mm^2^, the transient response of Voltron1 was larger than that of Voltron2. Thus, Voltron1 may outperform Voltron2 at spike detection under moderate illumination intensity.

We next sought to determine the kinetics with which voltage sensitivity increased under bright illumination and decreased under dim illumination. To measure these parameters, in HEK cells expressing either Voltron1_608_ or Voltron2_549_, we alternately clamped the voltage at −70 mV and +30 mV, and at each voltage we applied pulses of bright light (594 nm for JF_608_, 561 nm for JF_549_; 75 ms, 15 mW/mm^2^) interleaved with dim light (675 ms, 0.75 mW/mm^2^; Fig. 2c). By comparing the fluorescence at the two voltages during the dim-to-bright and bright-to-dim transitions, we measured the onset and decay of voltage sensitivity (Fig. 2d, e). For Voltron1_608_, voltage sensitivity arose with a time-constant of 5 ms and decayed with a time-constant of 46 ms (Fig. 2d). For Voltron2_549_, voltage sensitivity arose with a time-constant of 7 ms and decayed with a time-constant of 18 ms (Fig. 2e).

The appearance of the “normal” (i.e. previously reported) F vs. V behavior only at high illumination intensities, along with the finite time constants for voltage sensitivity to arise in response to a step-wise increase in illumination intensity, suggested that canonical Voltron voltage sensitivity might involve a photocycle intermediate, not the ground state (Fig 3). Sufficient 1P illumination populates the voltage-sensitive state, which then thermally relaxes to the dark-adapted state.

**Figure 3:**
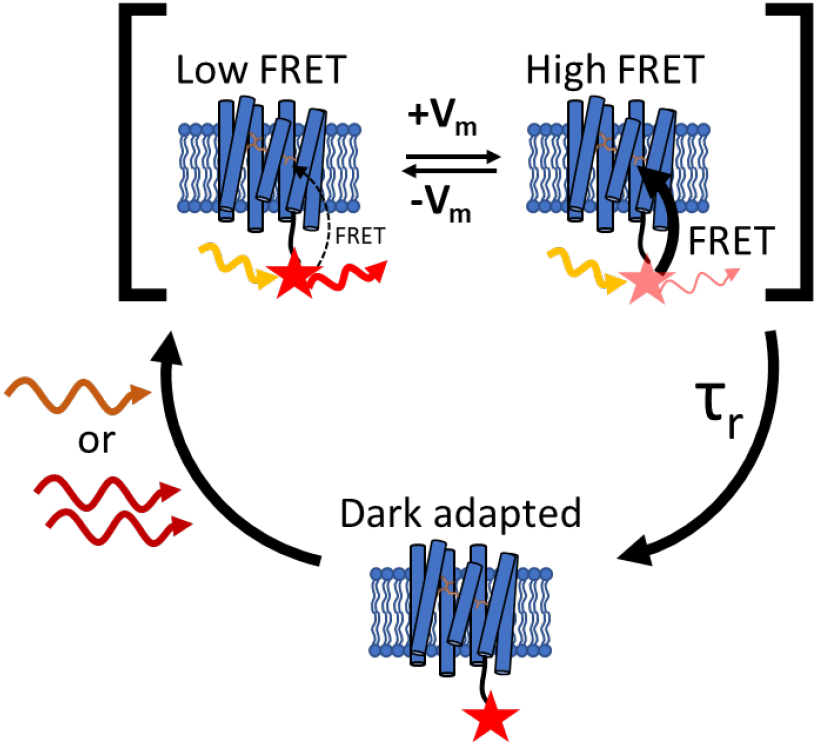
Model of photoactivated voltage sensitivity in a FRET-opsin voltage indicator. Dark-adapted Voltron2 does not show voltage-dependent FRET. Upon absorption of at least one photon with the proper energy (e.g. 594 nm or 1135 nm 2P), the opsin enters a voltage-sensitive equilibrium between high and low FRET states. The voltage sensitive equilibrium relaxes to the dark-adapted state with a time constant τ_r_ ∼ 18 ms. Voltage imaging requires concurrent measurement of the FRET donor fluorescence and optical pumping of the voltage-sensitive photocycle intermediate.

### Two photon photophysics

We hypothesized that the previously reported poor 2P voltage sensitivity in opsin-based GEVIs^27,39^ might arise from failure to populate voltage-sensitive photocycle intermediates (Fig. 3). This hypothesis is consistent with our observations that sufficient 1P illumination was required to observe 1P voltage sensitivity and with the much lower per-molecule excitation rate of 2P vs. 1P excitation^39^.

JF_549_ was the first dye reported for use with Voltron^21^ and has a 2P excitation peak in the center of the 1030-1080 nm mid-IR window reachable by multiple femtosecond laser technologies. We expressed Voltron2_549_ in HEK cells, clamped the voltage at −70 mV, and applied voltage steps to −90 mV to +50 mV. We first imaged the sample with donut-scanned 1040 nm 2P excitation (9.4 mW, 1 kHz scan repetition rate), and then applied the same voltage steps immediately afterwards under widefield 1P excitation (26 mW/mm^2^, 532 nm). Under 2P illumination, the fractional voltage sensitivity was much smaller and the step-response kinetics much slower compared to 1P illumination of the same cell (Fig. 4a). At 9.4 mW/cell, the dye bleached with a time constant of 3 s (Fig. 4a, inset). The 2P voltage sensitivity was −0.036 ± 0.015 per 100 mV (n = 6 cells, mean ± s.e.m., Fig 4d,e), significantly smaller than the 1P sensitivity of 0.10 ± .02 per 100 mV (n = 7 cells, mean ± s.e.m., Fig 4e) from paired measurements on the same set of cells (p = 0.04; two-tailed t-test, one cell only had a 1P recording; Fig. 4e).

**Figure 4:**
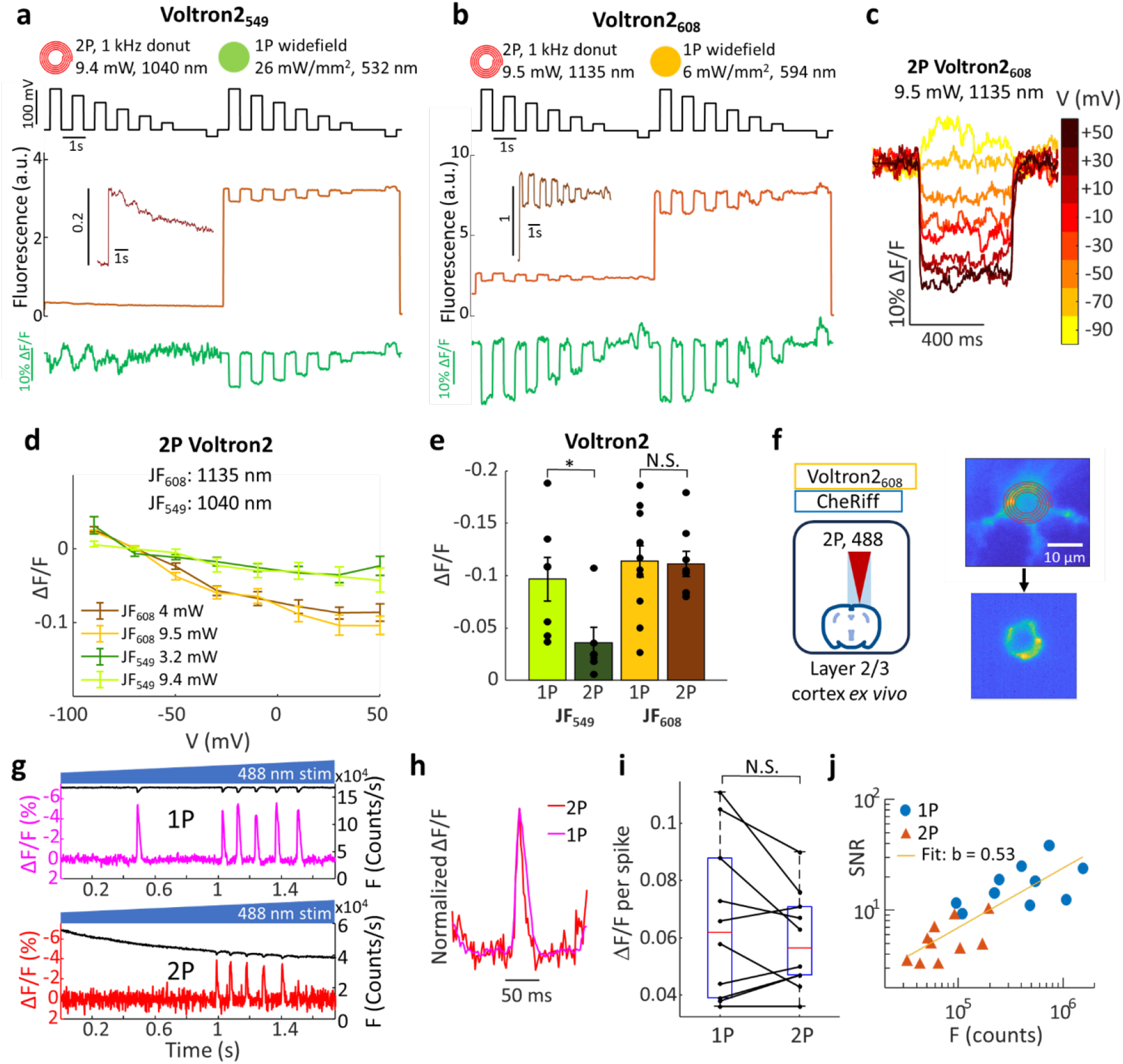
Two-photon voltage imaging with Voltron2. a) Comparison of Voltron2_549_ voltage sensitivity under 2P vs. 1P illumination. A HEK-293T cell expressing Voltron2_549_ was subjected to voltage steps from a holding potential of −70 mV. Fluorescence was recorded under 2P excitation and then under 1P excitation. The fractional changes in fluorescence were smaller and slower under 2P vs. 1P excitation. Inset shows magnified 2P fluorescence trace. Photobleaching was corrected before calculating ΔF/F. b) Same as (a), but with Voltron2_608_. c) Voltage step response from the same Voltron2_608_-expressing cell shown in panel (b). d) Voltage response under 2P excitation for Voltron2_549_ and Voltron2_608_, under dim (3-4 mW, dark colors) and bright (9.4-9.5 mW, light colors) illumination. Error bars s.e.m., *n* = 6-12 cells. e) Comparison of responses to 100 mV step (−90 to +10 mV) for Voltron2_549_ and Voltron2_608_ from matched samples and measurement conditions. Voltron2_549_ 1P: 26 mW/mm^2^, 532 nm, *n* = 7 cells; 2P: 9.4 mW, 1040 nm, *n* = 6 cells. Voltron2_608_ 1P: 6 mW/mm^2^, 594 nm, *n* = 11 cells; 2P: 9.5 mW, 1135 nm, *n* = 8 cells. Error bars mean ± s.e.m. 1P voltage sensitivity was significantly greater than 2P voltage sensitivity for Voltron2_549_ (p = 0.04 two-tailed t-test), but not for Voltron2_608_. f-j) 2P voltage imaging in acute brain slices co-expressing Voltron2_608_ and CheRiff. f) Left: experimental protocol. Right: (top) once a neuron was located, a 2P donut scan was targeted to the soma (500 Hz, 5.5 mW, 1135 nm.). (bottom) Fluorescence image recorded on a camera with donut-scan 2P excitation. g) Fluorescence traces (left axis, ΔF/F; right axis, F) of from 1P (top) and 2P (bottom) epochs of a single recording. A ramp of blue light from 0 to 0.5 mW/mm^2^ evoked spikes. h) Spike-triggered average traces from (g) were normalized and overlayed, demonstrating similar response kinetics for 1P and 2P recordings. i) Spike heights under matched 1P and 2P recordings were not significantly different (*n* = 10 cells, paired t-test). j) SNR vs. fluorescence (counts/cell/frame) for each recorded cell (*n* = 10 cells). The SNR and fluorescence show a power-law relationship with exponent b = 0.53 (95% confidence bounds: 0.35-0.72, R^2^ = 0.68), consistent with shot-noise limited SNR (b = 0.5) with the same relative signal level for 1P and 2P excitation. The higher SNR of the 1P recording can be attributed to the brighter fluorescence.

The 2P action spectrum for Voltron sensitization is not known, but prior work on 2P excitation of bacteriorhodopsin provides some guidance. For bacteriorhodopsin, 2P excitation of the S_1_ first excited state peaks at 1140 nm, whereas 2P excitation to the S_2_ second excited state (a symmetry-forbidden 1P transition) is a broad transition centered just under 1000 nm.^40^ Retinal isomerization and initiation of the photocycle require excitation to S_1_, so we reasoned that 2P excitation to S_1_ might favor voltage sensitivity while excitation to S_2_ might be unproductive or even counteract sensitization. This reasoning suggested that 2P excitation around 1140 nm might favor population of voltage sensitive states. JF_608_ dye shows 2P excitation with a peak around 1135 nm,^41^ so we reasoned that in Voltron2_608_, 2P light at 1135 nm might drive opsin sensitization and simultaneously excite the FRET donor for voltage imaging.

We expressed Voltron2_608_ in HEK cells, clamped the voltage at −70 mV, and applied voltage steps to - 90 mV to +50 mV under either bright (9.5 mW) or moderate (4 mW) 2P excitation at 1135 nm (Fig. 4b-e). The laser scan traced the periphery of the cell membrane at 1000 Hz. We then repeated the voltage steps under moderate (6 mW/mm^2^) 1P excitation at 594 nm (Fig. 4b). Under 2P excitation, Voltron2_608_ showed much larger voltage sensitivity than Voltron2_549_ (Fig. 4c-e), consistent with our hypothesis that long-wavelength 2P excitation was more effective at populating the voltage-sensitive photocycle intermediate.

For both Voltron2_549_ and Voltron2_608_, the voltage sensitivity under lower (3.2 - 4 mW) 2P power was similar to the sensitivity at 9.4 – 9.5 mW (Fig. 4d). This observation suggests that increasing 2P power beyond a few mW/cell would not increase the voltage sensitivity further, though the increase in overall brightness at higher laser power would increase the shot noise-limited SNR. Due to the low fluorescence signal under 2P conditions, we were not able to explore lower 2P powers.

We next explored 2P voltage imaging with Voltron2_608_ in mouse acute brain slices. We co-expressed Voltron2 and CheRiff, a blue light-activated channelrhodopsin, in layer 2/3 cortical pyramidal neurons via *in utero* electroporation. We prepared acute brain slices and incubated the slices with JF_608_ dye. We sequentially performed 1P (594 nm, 22 mW/mm^2^) and 2P (1135 nm, 25 mW) imaging on the same cells while evoking neuronal spikes with dim ramping (0-0.5 mW/mm^2^) 488 nm illumination (Fig. 4g). The spike-triggered average waveforms under 1P and 2P excitation matched closely, demonstrating that 1P and 2P recordings with Voltron2_608_ have the same time resolution.

We performed the same paired 2P and 1P recording protocol on pyramidal cells from cortical layer 2/3 and from hippocampus CA1. We varied 1P excitation power (22 - 70 mW/mm^2^) and 2P power (5.5 - 15 mW), to explore the effect of brightness in the sensitivity-saturated regime. Across 10 recordings (two recordings from each of 5 cells), we found no significant difference between 2P and 1P voltage sensitivity, ΔF/F per spike (Fig. 4i). For each recording, we also calculated the SNR (ratio of spike height to baseline noise) and plotted this against the fluorescence counts per cell per frame (Fig. 4j). On a log-log plot, the 2P and 1P data fell along the same line, with a slope of 0.53 (95% confidence bounds: 0.35-0.72, R^2^ = 0.68). This observation is consistent with both signals being shot-noise limited (slope = 0.5) with the same signal strength. Thus the greater SNR of the 1P recordings was attributed almost entirely to the increased fluorescence brightness under 1P conditions.

Finally, we tested 2P voltage imaging of Voltron2_608_ *in vivo*. We injected a viral vector for cre-dependent bicistronic expression of Voltron2 and CheRiff into Layer 1 barrel cortex of *Ndnf*-*Cre* mice. We performed optical stimulation and voltage imaging of layer 1 interneurons through a surgically implanted window in an anesthetized mouse. We sequentially performed 1P (594 nm, 30 mW/mm^2^) and 2P (1135 nm, 11-20 mW) imaging on the same cells while evoking neuronal spikes with pulses of 488 nm illumination of successively greater intensity (0.2-0.35 mW/mm^2^; Fig. 5a). The spike-triggered average spike waveforms under 1P and 2P excitation matched closely, displaying identical millisecond-scale spike kinetics.

**Figure 5:**
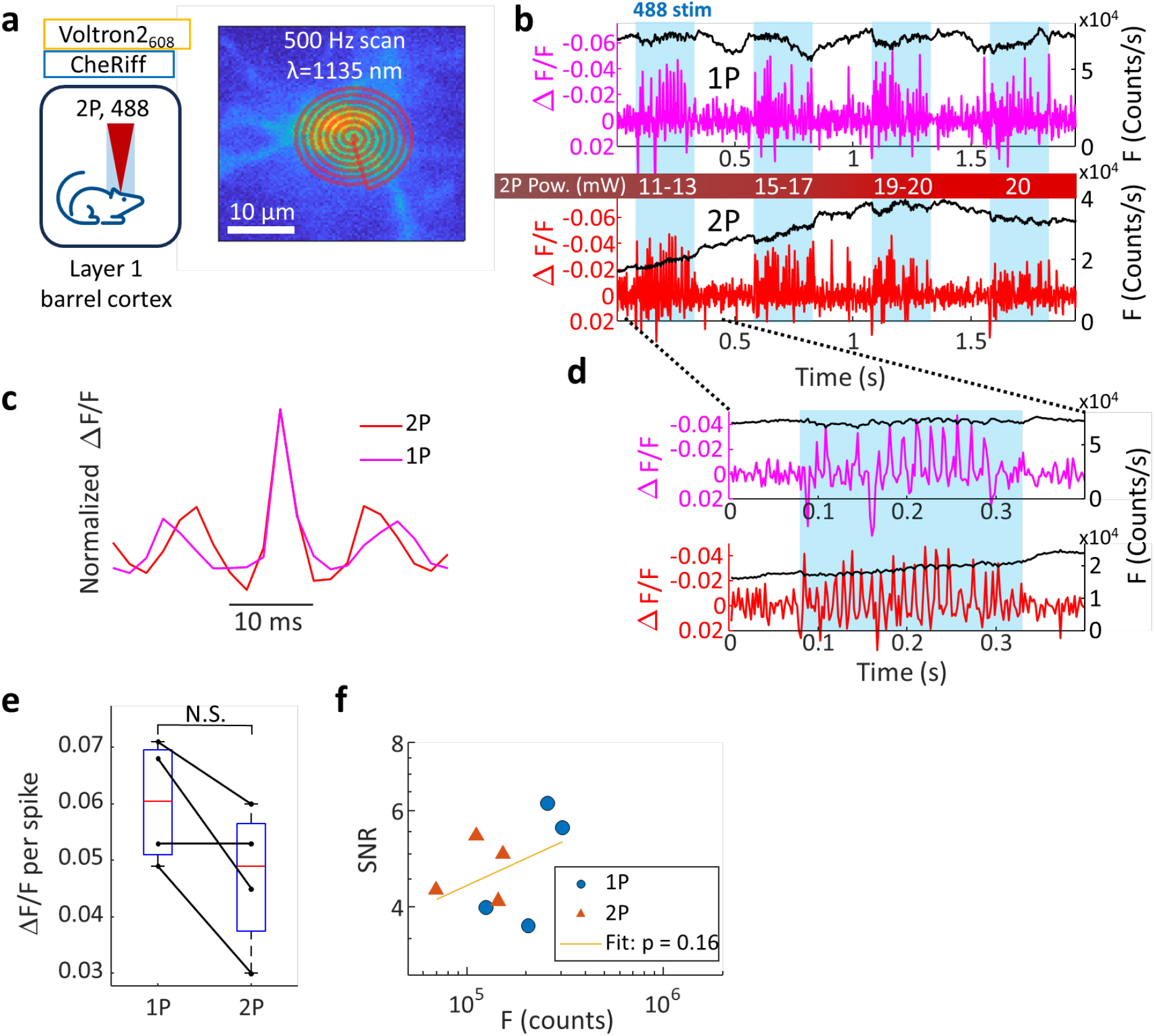
Two-photon voltage imaging with Voltron2_608_ *in vivo*. a) Cre-dependent Voltron2 and CheRiff were expressed by AAV injection in Layer 1 barrel cortex of *Ndnf-Cre* mice. JF_608_ dye was injected the day prior to the experiment, and neurons were imaged through a craniotomy under a 2P donut scan, 1135 nm, 500 Hz. b) Fluorescence traces (left axis, detrended ΔF/F; right axis, F) of from 1P (top) and 2P (bottom) epochs of a single recording. Pulses of blue light increasing from 0.2 to 0.35 mW/mm^2^ evoked spikes. 1P: 30 mW/mm^2^ 594 nm excitation; 2P: 1135 nm excitation at slowly increasing power to counteract photobleaching. Optogenetic stimulation evoked spikes which were detectable via voltage imaging. c) Spike-triggered average traces from (b) were normalized and overlayed, demonstrating similar response kinetics for 1P and 2P recordings (1P: N = 49 spikes; 2P: N = 54 spikes). d) Response to a single blue pulse. e) Spike heights under matched 1P and 2P recordings were not significantly different (*n* = 4 cells). f) SNR vs. fluorescence (counts/cell/frame) for each recorded cell (*n* = 4 cells, slope p = 0.16 ± 0.35 95% confidence interval; R^2^ = 0.18). Deviation from shot noise-limited SNR *in vivo* is likely due to contributions from background fluorescence, brain motion, and blood flow.

We compared statistics from four recordings (one from each of four cells). The 1P voltage sensitivity was better than the 2P sensitivity in three of the cells, but this did not reach statistical significance. The decreased 2P sensitivity may have been partially the result of bleaching at the higher 2P powers attained in these experiments. A log-log plot of SNR vs. fluorescence did not demonstrate a linear relationship (R^2^ = 0.18). For both 1P and 2P recordings, the noise had substantial contributions from background fluorescence, motion artefacts, and blood flow, leading to cell-specific noise above shot noise.

## Discussion

The absence of 2P voltage sensitivity with FRET-opsin reporters has long been a barrier in the field of voltage imaging. Here we show that 2P voltage imaging with FRET-opsin GEVIs is feasible if the illumination populates the voltage-sensitive photocycle intermediates. Achieving this goal required selecting a 2P excitation wavelength (1135 nm) that efficiently populated the intermediate state and a dye that was efficiently excited at this wavelength while also undergoing efficient FRET with the opsin, and applying scan patterns which revisited each molecule frequently enough to overcome relaxation of the voltage-sensitive intermediates. These results open the door to 2P voltage imaging *in vivo* with FRET-opsin GEVIs.

Our spectroscopic studies point to strategies for rational improvement of both 1P and 2P FRET-opsin GEVIs and voltage imaging systems. For instance, there may be other dyes whose excitation peak better matches the peak of the 2P opsin sensitization spectrum while still engaging in productive FRET with the voltage-sensitive states. Protein engineering efforts to slow the kinetics of relaxation of the voltagesensitive states could also enhance voltage sensitivity. Furthermore, one might engineer imaging systems in which 1P and 2P illumination are interleaved, with the 1P driving population of the voltagesensitive states and the 2P providing the excitation for optically sectioned imaging.

2P voltage imaging still faces difficulties as a practical tool for *in vivo* neural imaging. We recently compared the power budgets of 1P and 2P excitation: to achieve useful count rates for voltage imaging with a standard 80 MHz source, 2P excitation requires ∼10^4^-fold greater power per cell compared to 1P excitation.^39^ The maximum biologically safe laser power for 2P voltage imaging can be set by either average or peak illumination intensity. The time- and space-average power into the sample should be capped to avoid temperature rises greater than a few °C (5 °C can induce permanent damage, but smaller temperature rises may alter neural firing patterns).^42^ The peak intensity at the laser focus should not exceed ∼1 nJ/pulse, the saturation intensity of most fluorophores.^43^ At higher intensities, nonlinear photodamage might occur.

Our brightest 2P voltage recording was obtained at a power of just under 10 mW (0.125 nJ/pulse at 80 MHz), which induced bleaching with a time constant as short as 2.5 s (with scan-to-scan variation). Increases in peak pulse energy, offset by decreases in laser repetition rate, may enable brighter overall fluorescence signals and improvements in SNR, provided that the thermal and peak-intensity limits are respected. We discuss the optical and molecular constraints on 2P voltage imaging in detail in Ref. 39.

Our results also have important implications for use of Voltron2 under 1P excitation. For very long-term recordings, a natural inclination is to decrease the illumination intensity to avoid photobleaching or phototoxicity. However, our results show that this strategy may unintentionally lead to a loss of voltage sensitivity. A better strategy would be to interleave epochs of intense (> 10 mW/mm^2^) illumination with epochs of darkness. Similarly, for voltage imaging of large samples (e.g. an entire mouse heart), the excitation intensities may be low, leading to a loss of voltage sensitivity. To preserve sensitivity, one should either make an array of focal spots, or apply intermittent high-intensity illumination.

The complex photophysics of the FRET-opsin GEVIs suggest that future protein engineering efforts should be accompanied, at a minimum, by a quantification of intensity-dependent voltage sensitivity. An interesting avenue for future explorations would be to determine the photocycle basis for the intensity-dependent changes in voltage sensitivity and voltage step-response waveforms shown in Figs. 1 and 2.

## Materials and Methods

### Genetic constructs

Voltron1 and Voltron2 plasmids were obtained from Addgene (#119033 and #172909, respectively). For lentiviral transduction, the Voltron sequence was cloned into a lentiviral backbone with a CMV promoter using standard Gibson Assembly. Briefly, the vector was linearized by double digestion using restriction enzymes (New England Biolabs). DNA fragments were generated by PCR amplification and then fused with the backbones using NEBuilder HiFi DNA assembly kit (New England Biolabs). Resulting plasmids were verified by sequencing (GeneWiz).

For experiments in neurons, we co-expressed Voltron2 with a blue-shifted channelrhodopsin, CheRiff by a self-cleaving p2a linker. For *ex vivo* slice experiments, we used a previously-published whole-cell-expressing construct^10^ (Addgene #203228). For *in vivo* experiments, we generated a plasmid with soma-localized Voltron2 and soma-localized CheRiff under the hSyn promoter and flanked by LoxP sites for Cre recombinase-dependent expression. The genes were cloned into an adeno-associated virus (AAV) backbone using standard Gibson Assembly. AAV was produced by UNC Neurotools using the supplied plasmids.

### HEK cell culture

HEK293T cells were maintained in tissue culture-treated culture dishes (Corning) at 37 °C, 5% CO_2_ in Dulbecco’s Modified Eagle Medium supplemented with 10% fetal bovine serum, 1% GlutaMax-I, penicillin (100 U/mL), streptomycin (100 mg/mL). For each imaging experiment, cells in one 35 mm dish were either transiently transfected with the construct to be imaged using TransIT-293 lipofection reagent (Mirus Bio) or were virally transduced with a lentivirus. We saw no difference in voltage sensitivity or photophysics between the lipofected vs. virally transduced HEK cells. For lipofection, the construct was diluted 1:5 with empty pUC19 vector (New England Biolabs) and then transfected with 7.5 µL of TransIT-293 and 2.5 µg of DNA. Cells were replated 36-60 hours after transfection on glass-bottomed dishes (Cellvis, Cat. # D35-14-1.5-N) that were previously coated in poly-D-lysine to aid in cell adhesion.

### Lentiviral transduction

All the lentivirus preparations were made in house. HEK293T cells were co-transfected with the second-generation packaging plasmid psPAX2 (Addgene #12260), envelope plasmid VSV-G (Addgene #12259) and transfer plasmids at a ratio of 9:4:14. For small batches, 5.6 μg total plasmids for a small culture (300k cells in 35 mm dish) gave sufficient yield of lentivirus. Lentivirus was not further concentrated. For lentiviral transduction, 100 μL of lentivirus were added to a single 35 mm dish. After 48-60 hours, cells were either replated onto glass for imaging or split and replated on 35 mm plastic dishes for continued growth. Virally transduced cultures could be used for up to three passages after transduction. For all experiments, imaging was performed 12-24 hours after replating on glass.

### Electrophysiology and Buffers

Half an hour prior to imaging, the appropriate JF-HaloTag ligand dye was added to the medium in each dish of cells to a final concentration of 100 nM. Immediately prior to imaging, the medium was removed, and the cells were rinsed, then covered with dye-free extracellular (XC) buffer. The XC buffer contained 125 mM NaCl, 2.5 mM KCl, 3 mM CaCl_2_, 1mM MgCl_2_, 15 mM HEPES, 20 mM glucose, which was adjusted with NaOH to a pH of 7.3 and with sucrose to an osmolality of 305-310 mOsm, as measured by a vapor-pressure osmometer (Wescor). Filamented patch pipettes were pulled using an automated puller (Sutter P-1000) to a tip resistance of ∼6 MΩ and were filled with an intracellular buffer (IC) containing (in mM) 6 NaCl, 130 K-aspartate, 2 MgCl_2_, 5 CaCl_2_, 11 EGTA, and 10 HEPES, with pH adjusted to 7.2 by KOH^44^.

Whole-cell voltage clamp was acquired using a modified syringe to manipulate pressure, following Li.^44^

### Microscope and illumination control

Single-photon (1P) imaging experiments were performed on a custom-built inverted microscope with a computer-controlled patch amplifier (Axon Instruments, Multiclamp 700B). Once a whole-cell patch was established, acquisition was controlled using custom MATLAB/C++ acquisition software (https://www.luminosmicroscopy.com/). The illumination path contained a 594 nm laser (Hübner Photonics, Cobolt Mambo) and a 561 nm laser (Hübner Photonics, Cobolt Jive). The laser outputs were modulated using a multichannel acousto-optic tunable filter (Gooch & Housego, TF525-250-6-3-GH18A with MSD040-150 driver), and imaging was performed through a high-NA 60x water-immersion objective (Olympus UPLSAPO60XW, 0.28 mm working distance, NA = 1.2) onto an sCMOS camera (Hamamatsu, ORCA-Flash 4.0). Imaging of JF_549_ and JF_608_ was performed through a 488/561/633 nm triband dichroic (Chroma) and a 405/488/594 nm triband dichroic (Semrock), respectively. A 594 nm long-pass emission filter was used for both dyes (Semrock, BLP01-594R-25). Electrical waveforms and measurements were transduced through a computer-controlled data acquisition device (National Instruments, PCIe-6343).

The sample was placed on a 2-axis motorized stage (Ludl Electronic Products, MAC6000), and a 3-axis micromanipulator was used for patch pipette control (Sutter, MP-285).

Two-photon (2P) imaging experiments were performed on a custom-built upright microscope equipped with 1P and 2P illumination paths, a shared emission path to an sCMOS camera (Hamamatsu, ORCA-Flash 4.0), and a computer-controlled patch amplifier (Axon Instruments, Axopatch 200B). An 80 MHz tunable ultrafast laser (Spectra-Physics, InSight DeepSee) was modulated using an electro-optic modulator (ConOptics, 350-80-02 with 302RM driver) and directed using a pair of galvanometric mirrors (Cambridge Technologies 6215H with 671HP driver). 488 nm (Coherent OBIS 488-100 LS), 532 nm (Laserglow LLS-05320PFM-00159-01), and 594 nm (Hübner Photonics Cobolt Mambo 0594-04-01-0100-500) lasers were combined and independently modulated using a multichannel acousto-optic tunable filter (Gooch & Housego PCAOM NI-VIS with MSD040-150 driver). The 1P lasers were patterned using a digital micromirror device (Vialux V-7001). Imaging of HEK cells and brain slices was performed through a high-NA 25x water-immersion objective (Olympus XLPLN25XWMP2, 2 mm working distance, NA=1.05).

*In vivo* imaging was performed through a high-NA 25x water-immersion objective (Olympus XLPLN25XSVMP2, 4 mm working distance, NA = 1). A 785 nm long-pass dichroic (Semrock Di03-r785-t3) separated the 2P excitation from the 1P and imaging paths. A 594 nm long-pass dichroic (Semrock Di03-r594-t3) separated the 1P excitation light from the imaging path. Emission filters at 628/40 and 593/40 were used for imaging of JF_608_ and JF_549_, respectively. Electrical waveforms and measurements were transduced through a computer-controlled data acquisition device (National Instruments, PCIe-6363).

Galvo control and feedback waveforms were transduced through a second computer-controlled data acquisition device (National Instruments, PCIe-6343). The sample was placed on a motorized 2-axis state, with focus controlled by objective displacement and a 3-axis micromanipulator used for patch pipette control (Sutter MPC-200 controller with MPC-78 stage and MP-285 manipulators).

Camera scaling was calibrated using a stage micrometer (Thorlabs, R1L3S2P), and illumination powers were calibrated using a power meter (Thorlabs, PM400) with either a photodiode (Thorlabs, S170C) or a thermal (Thorlabs, S175C) slide power sensor, for 1P and 2P, respectively.

### Animals

All animal procedures adhered to the National Institutes of Health Guide for the care and use of laboratory animals and were approved by the Harvard University Institutional Animal Care and Use Committee (IACUC).

### In utero electroporation (IUE)

The IUE surgery was performed as described previously.^9^ Timed-pregnant female CD1 mice (embryonic day 15.5, E15.5; Charles River) were deeply anesthetized and maintained with 2% isoflurane. The animal body temperature was maintained at 37 °C. Uterine horns were exposed and periodically rinsed with warm phosphate-buffered saline (PBS). Plasmid DNA was diluted in PBS (2 μg/μL; 0.05% fast green), and 1 µL of the mixture was injected into the left lateral ventricle of the embryos. Electrical pulses (40 V, 50 ms duration) targeting the hippocampus were delivered five times at 1 Hz using tweezers electroporation electrodes (CUY650P5; Nepa Gene). Injected embryos were returned to the abdominal cavity, and the surgical incision was closed with absorbable PGCL25 sutures (Patterson).

### Cranial window surgery

Surgeries were conducted on NDNF-cre X NPY-GFP mice of both sexes, following the protocol outlined by ^45^. The surgical procedure began by exposing the skull, followed by a 3 mm circular craniotomy at coordinates 3.3 – 3.4 mm lateral and 1.6 caudal relative to the bregma. The craniotomy was made using a dental drill. Following this, a stack of one 5 mm and two 3 mm round cover glass (Thomas Scientific, 1217N66), pre-glued by optical glue (Norland 61), was inserted into the opening. All subsequent experimental procedures were carried out at least one week post-surgery, ensuring that the health of each mouse was stable.

### AAV injection

Following craniotomy, AAV virus (final titer ∼4e12 GC/mL) was injected at a rate of 45 nL/min using a home-pulled micropipette (Sutter P1000 pipette puller) mounted in a microinjection pump (World Precision Instruments Nanoliter 2010) controlled by a micro-syringe pump controller (World Precision Instruments Micro4). The micropipette was positioned using a stereotaxic instrument (Sutter Instruments).

### In Vivo Imaging

JF_608_-HaloTag ligand solution was first prepared as previously described^41^, and was then retro-orbitally delivered 24 hours before the imaging session. Mice were lightly anaesthetized (0.7-1% isoflurane) and head-fixed under the upright microscope using a titanium head plate. Eyes were kept moist using ophthalmic eye ointment. The body temperature was continuously monitored and maintained at 37°C using a heating pad (WPI, ATC-2000). A typical imaging session lasted 2-3 hours, after which the animals quickly recovered and were returned to their home cage.

### Brain slice preparation

Coronal slices (300 µm) were prepared from CD1 mice of either sex between 2-4 postnatal weeks. Animals were anesthetized with isoflurane and euthanized by decapitation. The brain was then removed and placed in ice-chilled slicing solution containing (in mM): 210 sucrose, 3 KCl, 26 NaHCO_3_, 1.25 NaH_2_PO_4_, 5 MgCl_2_, 10 D-glucose, 3 sodium ascorbate, and 0.5 CaCl_2_ (saturated with 95% O_2_ and 5% CO_2_). Acute slices were made using a Vibratome (VT1200S, Leica) while maintained in the slicing solution.

Slices were recovered at 34 °C for 10 min in the imaging solution (artificial cerebrospinal fluid, ACSF) containing (in mM): 124 NaCl, 3 KCl, 26 NaHCO_3_, 1.25 NaH_2_PO_4_, 2 MgCl_2_, 15 D-glucose, and 2 CaCl_2_ (saturated with 95% O_2_ and 5% CO_2_). Slices were then incubated in ACSF containing JF_608_-HaloTag ligand^46^ (0.5-1 µM) for 30 min at room temperature, and moved to a fresh ACSF for another 30 min to wash out excess dye. Slices were maintained and recorded at room temperature, 24 °C.

### Analysis

All analyses were performed using custom semi-automated MATLAB code. All automated analysis pipelines included a quality-control checkpoint at which key fits and values were manually checked. Analysis of 1P-only experiments began with manual selection of a membrane ROI from each cell.

Analysis of experiments including 2P excitation began with automated selection of an ROI based on the 2P fluorescent signal. In both cases, the same ROI was used for each experiment from a given cell.

Analysis proceeded by extracting a time trace of fluorescence by equal-weighted averaging of photon counts over the selected ROI followed by subtraction of the mean counts from a background region selected near the cell. The resulting one-dimensional time trace of fluorescence was used for all subsequent analyses. This trace was smoothed with a moving mean filter of window 20 ms for display of single voltage step responses and a moving mean filter of window 10 ms for calculation of the ΔF/F traces in Fig. 1l and Fig. 2d,e. Photobleaching was corrected using a bi-exponential fit to each illumination epoch. This fit provided the baseline from which values of ΔF were calculated, and which was used for normalization of ΔF/F.

Neural spike recordings were detrended by smoothing the data with a moving mean filter of width 50 ms. This produced a smooth baseline from which ΔF/F could be calculated for the raw fluorescence trace. Spike heights, noise level, and fluorescence levels were manually extracted from all recordings before plotting. The logarithm of the SNR and per-cell counts were taken, and a linear fit was performed using MATLAB’s curve fitting toolbox.

Significance of paired conditions was calculated using a two-tailed paired sample t-test, and significance of unpaired data was calculated using a two-tailed two-sample t-test, both using MATLAB’s machine learning and statistics toolbox.

## Acknowledgements

We thank Daniel Itkis, Andrew Preecha, Shahinoor Begum, David Wong-Campos, Byung Hun Lee, and Ethan Perets for technical assistance and helpful discussions. This work was supported by NIH grants 1R01NS133755, 1RF1NS126043, and NSF Quantum Sensing for Biophysics and Bioengineering (QuBBE) Quantum Leap Challenge Institute (QLCI) grant OMA-2121044.

F.P.B., H.C.D, and A.E.C. designed the study and experiments. F.P.B. and A.E.C. wrote the manuscript with input from all authors. F.P.B. collected the single-photon data. F.P.B. and H.C.D. collected the two-photon HEK-293 data. P.P. prepared the I.U.E construct, performed I.U.E, prepared brain slices, and assisted F.P.B. with data collection for 2P acute slice imaging. Y.Q. prepared the AAV construct, conducted virus injection and cranial window surgery and assisted F.P.B. with data collection for 2P *in vivo* imaging. F.P.B. performed all analyses with critical input from H.C.D. and A.E.C. A.E.C. supervised the work.

The authors declare that they have no competing interests.

Voltage imaging data will be made available on the DANDI archive.

## References

1. Adam, Y. All-optical electrophysiology in behaving animals. Journal of Neuroscience Methods 353, 109101 (2021).

2. Abdelfattah, A. S. et al. Sensitivity optimization of a rhodopsin-based fluorescent voltage indicator. Neuron 111, 1547-1563.e9 (2023).

3. Böhm, U. L. et al. Voltage imaging identifies spinal circuits that modulate locomotor adaptation in zebrafish. Neuron 110, 1211-1222.e4 (2022).

4. Adam, Y. et al. Voltage imaging and optogenetics reveal behaviour-dependent changes in hippocampal dynamics. Nature 569, 413 (2019).

5. Kralj, J. M., Hochbaum, D. R., Douglass, A. D. & Cohen, A. E. Electrical spiking in Escherichia coli probed with a fluorescent voltage indicating protein. Science 333, 345–348 (2011).

6. Akemann, W. et al. Imaging neural circuit dynamics with a voltage-sensitive fluorescent protein. J.Neurophysiol. 108, 2323–2337 (2012).

7. Gong, Y. et al. High-speed recording of neural spikes in awake mice and flies with a fluorescent voltage sensor. Science 350, 1361–1366 (2015).

8. Wong-Campos, J. D. et al. Voltage dynamics of dendritic integration and back-propagation in vivo. (2023) doi:10.1101/2023.05.25.542363.

9. Landau, A. T. et al. Dendritic branch structure compartmentalizes voltage-dependent calcium influx in cortical layer 2/3 pyramidal cells. eLife 11, e76993 (2022).

10. Park, P. et al. Dendritic voltage imaging reveals biophysical basis of associative plasticity rules. 2023.06.02.543490 Preprint at 10.1101/2023.06.02.543490 (2023).

11. Jia, B. Z., Qi, Y., Wong-Campos, J. D., Megason, S. G. & Cohen, A. E. A bioelectrical phase transition patterns the first vertebrate heartbeats. Nature 622, 149–155 (2023).

12. Sacconi, L. et al. KHz-rate volumetric voltage imaging of the whole Zebrafish heart. Biophysical Reports 2, 100046 (2022).

13. Azimi Hashemi, N. et al. Rhodopsin-based voltage imaging tools for use in muscles and neurons of Caenorhabditis elegans. Proceedings of the National Academy of Sciences 116, 17051–17060 (2019).

14. Kralj, J. M., Douglass, A. D., Hochbaum, D. R., Maclaurin, D. & Cohen, A. E. 3248630; Optical recording of action potentials in mammalian neurons using a microbial rhodopsin. Nat Methods 9, 90–95.

15. Piatkevich, K. D. et al. A robotic multidimensional directed evolution approach applied to fluorescent voltage reporters. Nature Chemical Biology 14, 352–360 (2018).

16. Kralj, J. M., Douglass, A. D., Hochbaum, D. R., Maclaurin, D. & Cohen, A. E. Optical recording of action potentials in mammalian neurons using a microbial rhodopsin. Nat. Meth. 9, 90–95 (2012).

17. Hochbaum, D. R. et al. All-optical electrophysiology in mammalian neurons using engineered microbial rhodopsins. Nat.Methods 11, 825–833 (2014).

18. Zou, P. et al. Bright and fast multicoloured voltage reporters via electrochromic FRET. Nature Communications 5, 4625 (2014).

19. Gong, Y., Wagner, M. J., Li, J. Z. & Schnitzer, M. J. Imaging neural spiking in brain tissue using FRET-opsin protein voltage sensors. Nat. Commun. 5, e3674 (2014).

20. Kannan, M. et al. Dual-polarity voltage imaging of the concurrent dynamics of multiple neuron types. Science 378, eabm8797 (2022).

21. Abdelfattah, A. S. et al. Bright and photostable chemigenetic indicators for extended in vivo voltage imaging. Science 365, 699–704 (2019).

22. Abdelfattah, A. S. et al. A general approach to engineer positive-going eFRET voltage indicators. Nat Commun 11, 3444 (2020).

23. Liu, S. et al. A far-red hybrid voltage indicator enabled by bioorthogonal engineering of rhodopsin on live neurons. Nat. Chem. 13, 472–479 (2021).

24. Chien, M.-P. et al. Photoactivated voltage imaging in tissue with an Archaerhodopsin-derived reporter. Science Advances 7, eabe3216 (2021).

25. Chamberland, S. et al. Fast two-photon imaging of subcellular voltage dynamics in neuronal tissue with genetically encoded indicators. Elife 6, e25690 (2017).

26. Liu, Z. et al. Sustained deep-tissue voltage recording using a fast indicator evolved for two-photon microscopy. Cell 185, 3408-3425.e29 (2022).

27. Brinks, D., Klein, A. J. & Cohen, A. E. Two-photon fluorescence lifetime imaging microscopy (2P-FLIM) of genetically encoded voltage indicators as a probe of absolute membrane voltage. Biophys. J. 109, 914–921 (2015).

28. Bando, Y., Sakamoto, M., Kim, S., Ayzenshtat, I. & Yuste, R. Comparative Evaluation of Genetically Encoded Voltage Indicators. Cell Reports 26, 802-813.e4 (2019).

29. Hendler, R. W., Shrager, R. I. & Bose, S. Theory and Procedures for Finding a Correct Kinetic Model for the Bacteriorhodopsin Photocycle. J. Phys. Chem. B 105, 3319–3328 (2001).

30. Lanyi, J. K. Bacteriorhodopsin. Annu. Rev. Physiol. 66, 665–688 (2004).

31. Bayraktar, H. et al. Ultrasensitive Measurements of Microbial Rhodopsin Photocycles Using Photochromic FRET. Photochemistry and Photobiology 88, 90–97 (2012).

32. Maclaurin, D., Venkatachalam, V., Lee, H. & Cohen, A. E. Mechanism of voltage-sensitive fluorescence in a microbial rhodopsin. Proceedings of the National Academy of Sciences 110, 5939– 5944 (2013).

33. Kübel, J. et al. Transient IR spectroscopy identifies key interactions and unravels new intermediates in the photocycle of a bacterial phytochrome. Physical Chemistry Chemical Physics 22, 9195–9203 (2020).

34. Penzkofer, A., Silapetere, A. & Hegemann, P. Photocycle dynamics of the Archaerhodopsin 3 based fluorescent voltage sensor Archon2. Journal of Photochemistry and Photobiology B: Biology 225, 112331 (2021).

35. Silapetere, A. et al. QuasAr Odyssey: the origin of fluorescence and its voltage sensitivity in microbial rhodopsins. Nat Commun 13, 5501 (2022).

36. Penzkofer, A., Silapetere, A. & Hegemann, P. Theoretical investigation of the photocycle dynamics of the Archaerhodopsin 3 based fluorescent voltage sensor Archon2. Journal of Photochemistry and Photobiology A: Chemistry 437, 114366 (2023).

37. Venkatachalam, V. et al. Flash memory: photochemical imprinting of neuronal action potentials onto a microbial rhodopsin. J.Am.Chem.Soc. 136, 2529–2537 (2014).

38. Hou, J. H., Venkatachalam, V. & Cohen, A. E. Temporal dynamics of microbial rhodopsin fluorescence reports absolute membrane voltage. Biophys.J. 106, 639–648 (2014).

39. Davis, H. C., Brooks, F. P., Wong-Campos, J. D. & Cohen, A. E. Optical constraints on two-photon voltage imaging. 2023.11.18.567441 Preprint at 10.1101/2023.11.18.567441 (2023).

40. Birge, R. R. & Zhang, C. Two-photon double resonance spectroscopy of bacteriorhodopsin. Assignment of the electronic and dipolar properties of the low-lying 1A*-g-like and 1B*+u-like p, p* states. The Journal of Chemical Physics 92, 7178–7195 (1990).

41. Grimm, J. B. et al. A general method to fine-tune fluorophores for live-cell and in vivo imaging. Nature Methods 14, 987–994 (2017).

42. Podgorski, K. & Ranganathan, G. Brain heating induced by near-infrared lasers during multiphoton microscopy. J.Neurophysiol. 116, 1012–1023 (2016).

43. Charan, K., Li, B., Wang, M., Lin, C. P. & Xu, C. Fiber-based tunable repetition rate source for deep tissue two-photon fluorescence microscopy. Biomed Opt Express 9, 2304–2311 (2018).

44. Li, C. A reliable whole cell clamp technique. Advances in Physiology Education 32, 3 (2008).

45. Goldey, G. J. et al. Removable cranial windows for long-term imaging in awake mice. Nature Protocols 9, 2515-2538-2515–2538 (2014).

46. Lin, D. et al. Time-tagged ticker tapes for intracellular recordings. Nat Biotechnol 41, 631–639 (2023).

